# Large-scale reorganization of DNA methylation and upregulation of extracellular matrix genes in the dorsal dentate gyrus following cocaine taking

**DOI:** 10.64898/2026.03.02.708518

**Authors:** Madelyn R. Baker, Elizabeth Brindley, Kyle A. Windisch, Rose Sciortino, Anjali Rajadhyaksha, Miklos Toth

## Abstract

Cocaine self-administration induces neurobiological adaptations in brain circuits involved in encoding reward-associated context. The dorsal hippocampus, particularly the dorsal dentate gyrus, plays a critical role in the precise encoding of spatial and contextual information. We hypothesized that the dentate gyrus is uniquely positioned to undergo epigenomic and transcriptomic changes because of the convergence of the contextual features of reward and cocaine-enhanced dopamine and norepinephrine signals during volitional drug-taking. We report that cocaine self-administration produces significant DNA methylation changes at an unusually large number of ∼30,000 genomic regions (>10%, q<0.01) in dentate granule cells (DGCs) of male mice. Cocaine preferentially hypomethylated regions with heterogenous methylation, switching the methylation state in ∼16% of DGCs on average. The cocaine-sensitive/responsive epigenomic regions were overrepresented in enhancers and were associated with 9,833 genes, many of which were involved in diverse functions relevant to neuronal functioning. However, among the differentially methylated genes only two regulatory genes, *c-fos* and *cartpt* (known to be activated by cocaine), and a cluster of genes encoding components of the extracellular matrix (implicated in neuroplasticity) were differentially expressed (mostly upregulated) following cocaine self-administration, suggesting a gene regulatory network that is transcriptionally robust to perturbations but still specific for context-driven and reward associated neuroplasticity in DGCs. Overall, our data demonstrates that cocaine self-administration induces epigenomic and transcriptomic changes in the dorsal dentate gyrus that may contribute to dorsal hippocampal plasticity and contextual memory associated with cocaine self-administration.

## Introduction

Cocaine use disorder is a chronic brain disease associated with persistent drug-taking and long-lasting neurobiological adaptations. These adaptations are thought to be mediated by long-lasting molecular changes, including the epigenomic remodeling of gene expression, within reward– and context-related brain circuits^1, 2^. Methylation of DNA, most frequently at CpG dinucleotides, is a common epigenomic modification that has been shown to be malleable by cocaine. For example, cocaine sensitization induces DNA methylation changes in the frontostriatal addiction circuits^3^ and cocaine self-administration (SA) alters DNA methylation in the medial prefrontal cortex (mPFC)^4^. Further, pharmacological inhibition of DNA methylation in the mPFC was reported to increase cocaine SA, suggesting a causal relationship between drug-induced DNA methylation and drug-taking behaviors^5, 6^. Similarly, several studies have reported a correlation between DNA methylation in the nucleus accumbens (NAc), a region central to the rewarding properties of cocaine^7^, and cocaine-induced behaviors^8, 9^. Overexpression of the neuronal DNA methyltransferase Dnmt3a in the NAc attenuates cocaine SA while inhibition of DNA methylation potentiates cocaine SA^10^. Finally, dietary methyl supplementation attenuates cocaine-seeking behavior and cocaine-induced *c-fos* activation in a DNA methylation-dependent manner^6^.

In addition to the mesolimbic reward system, the dorsal hippocampus plays a key role in addiction-related behaviors, particularly in encoding the contextual aspects of the experience^11–19^. However, the molecular mechanisms underlying cocaine-induced plasticity in this brain region remains less explored. The dorsal dentate gyrus, a hippocampal subregion that serves as the primary input to the hippocampal circuit, receives glutamatergic inputs that carries contextual and spatial information encoded in the entorhinal cortex. The dorsal dentate gyrus also receives dopamine inputs from the ventral tegmental area^20^ and norepinephrine projections from the locus coeruleus, which additionally release dopamine^21^. The principal neurons in the dorsal dentate gyrus are dentate granule cells (DGCs), which are essential for the initial encoding and early consolidation of contextual memories^22^. Importantly, DGCs have been shown to generate the contextual memories associated with cocaine exposure in conditioned place preference^23^ and cocaine SA environment^24^. The dentate gyrus further gates information flow to the CA3 and connected CA1 neurons^25^, which in turn, project to the NAc, as well as to the mPFC and amygdala^26, 27^. Given the convergence of the contextual features of cocaine reward-related behaviors and the rich landscape of cocaine-enhanced neurotransmitter signaling^28–30^, we hypothesized that the dentate gyrus is uniquely positioned to undergo epigenomic and transcriptomic changes during drug-taking.

To begin to examine the impact of cocaine on DNA methylation in the dorsal dentate gyrus, we performed genome-wide DNA methylation profiling and bulk RNA sequencing with dorsal DGCs from mice following intravenous SA. We show that cocaine SA induces significant DNA methylation changes, predominantly hypomethylation, at an unusually large scale involving nearly 10,000 genes. These changes cluster within genomic regions enriched in transcriptional enhancers and are associated with functionally diverse but neuron-relevant genes. Notably, a group of genes, encoding extracellular matrix (ECM) proteins, emerged as differentially expressed, suggesting a potential direct link to cocaine-induced hippocampal plasticity.

## Materials and Methods

### Animals

Animal experiments were performed in accordance with The Rockefeller University and Weill Cornell Medicine Institutional Animal Care and Use Committee guidelines. Adult C57BL/6 male mice (11 weeks upon arrival; Jackson Laboratory, Bar Harbor, ME) were group housed in a climate-controlled environment, 4-5 animals per cage, with a reverse 12 h light/dark cycle (7PM-7AM). Food and water were available *ad libitum*.

### Surgery

Following one-week habituation, mice underwent jugular catheterization surgery as described previously^31, 32^.

### Intravenous self-administration

Mice had 7 consecutive daily 2-hour cocaine intravenous SA (n=12) or yoked saline (n=11) sessions. Mice were trained to nose poke at the active hole of the chamber for delivery of 0.5 mg/kg infusion of cocaine on a fixed ratio 1 (FR1) reinforcement schedule. For the yoked saline controls, animals received an equivalent infusion of saline when their yoked paired cocaine SA mouse received an infusion of cocaine. Three cocaine SA mice were excluded from analysis (two lost catheter patency and one failed to self-administer).

### DNA Methylation Profiling by eRRBS

Genomic DNA was isolated from microdissected dorsal dentate granule cell layers obtained from frozen brain sections and processed for enhanced reduced representation bisulfite sequencing (eRRBS) as reported previously^33^.

### Bulk RNA-sequencing

RNA was extracted from microdissected granule cell layers and sequenced using standard poly(A)-selected bulk RNA-seq. Reads were aligned to the mm10 mouse genome and quantified using established pipelines. Differential expression analysis was performed with DESeq2 after low-count filtering and outlier removal; significance was defined by adjusted *p* < 0.1 and |logl_FC| > logl_ (1.3).

### Statistical Analysis

Behavioral data were analyzed with repeated measures two-way ANOVA (nose poke type (active versus inactive) x session) followed by Tukey multiple comparison posthoc test when main effect was observed. Significance of overlaps between differentially expressed and differentially methylated genes was determined using hypergeometric testing (*phyper* in R). Statistics for eRRBS and RNA-Seq are incorporated into the supplementary method section. Sample sizes were based on previous work in the lab and are similar to other published work in the field. Sample sizes for individual experiments are indicated in figure legends.

## Results

### Large-scale reorganization of DNA methylation by cocaine self-administration in dorsal dentate granule cells

To determine if cocaine SA in mice alters DNA methylation in dorsal DGCs, male mice were trained on an FR1 schedule in two-hour daily sessions at 0.5 mg/kg/infusion of cocaine for seven days (**Figure 1A**). Animals made significantly more active nose pokes (that led to cocaine administration) compared to inactive nose pokes (that did not lead to cocaine administration) through the 7-day trial that was already significant on the first day indicating learning that active nose pokes trigger cocaine infusion and drug reward (**Figure 1B**). Similarly, high intake of cocaine was established from the first day of the trial (**Supplementary Figure 1**). There was no difference in nose poke responding on the two holes that had no programmed consequence.

**Figure 1.**
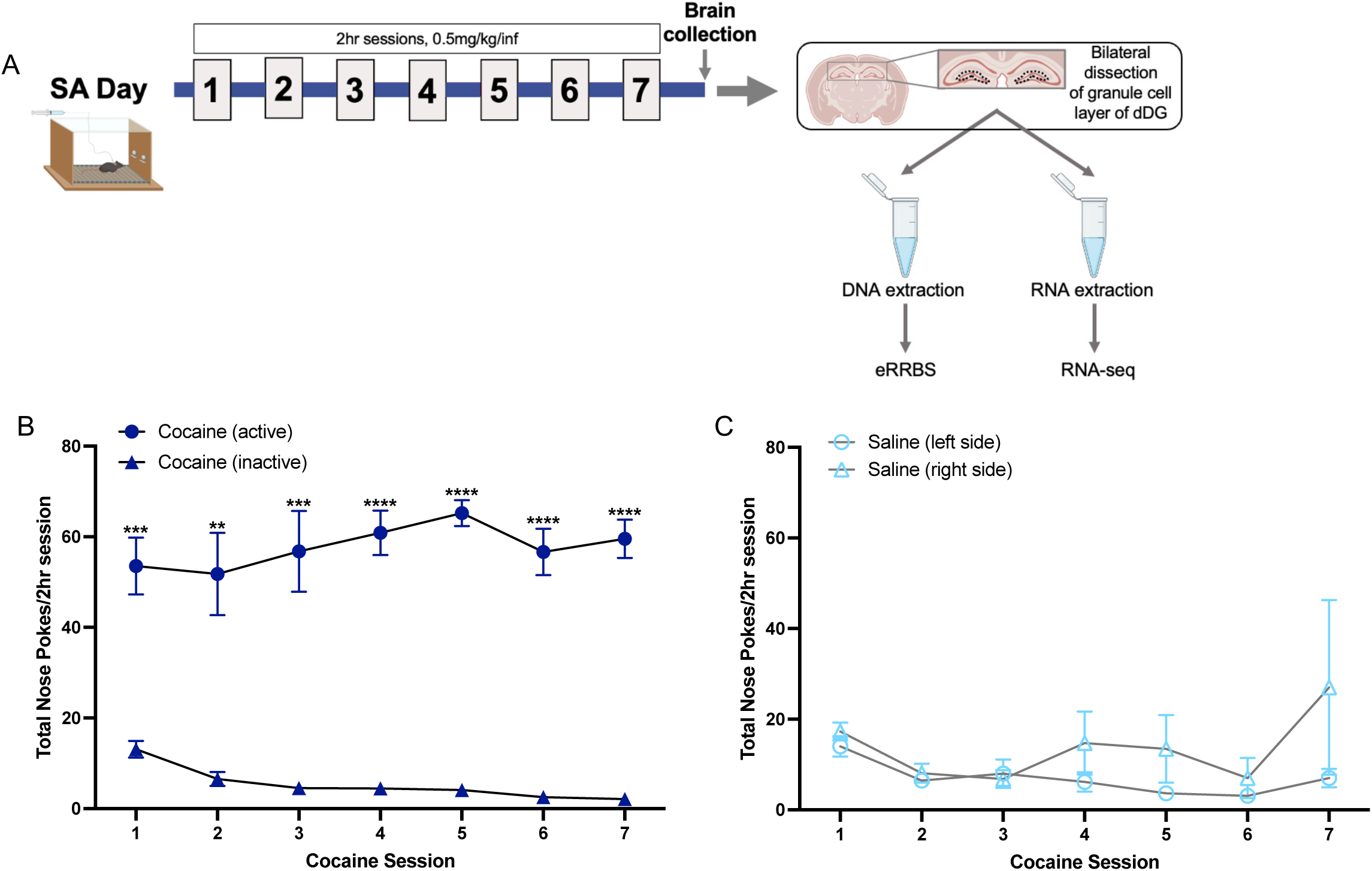
Experimental diagram of cocaine SA. (**A**) Experimental timeline of seven days of SA (2-hour sessions, 0.5 mg/kg/infusion), and representation of dissection of dorsal dentate gyrus DGC cell bodies and DNA isolation for eRRBS and RNA isolation for RNA-sequencing. (**B**) Number of nose pokes through seven sessions of cocaine SA. Two-way repeated measures ANOVA; main effect of poke type (F_1, 18_ = 151.60, p-value < 0.0001). Interaction between nose pokes and sessions (F_26, 108_ = 2.19, p-value = 0.049); Tukey multiple comparison test, **p=0.0011, ***p<0.0005, **** p < 0.0001. N= 12. (**C**) Yoked saline control animals did not show a significant difference in nose port pokes. Data are displayed as mean ± SEM. Two-way repeated measures ANOVA (poke type x session); no main effect of poke type (F_1,20_= 1.249; P = 0.2770) and no significant interaction (F_6,120_= 1.015; P = 0.4188). N=11. Data are displayed as mean ± SEM.

At the end of the 7-day SA trial, the granule cell layer was microdissected from dorsal hippocampal sections (by carefully separating it from the molecular layer and hilus) of cocaine and yoked control mice. Due to the laminar structure of the dentate gyrus, this region is comprised of the cell bodies of DGCs with minimal contamination from nuclei of other cell types^34, 35^. DNA methylation was profiled by eRRBS^33^ that is optimized for CpG-containing regions, including genomic regions with regulatory potential, as opposed to sequencing regions with no or few CpGs (**Figure 1A**). We identified 1,510,384 CpG sites that were covered across all samples (4 cocaine SA and 4 saline controls) by at least 10 reads (**Figure 2A**) and calculated methylation differences for each of these CpGs by subtracting the percent of copies methylated in yoked control DGCs from the percent of copies methylated in cocaine SA DGCs. Of these CpGs, 334,380 (22%) exhibited significant methylation differences (q-value ≤ 0.01) between SA and yoked control DGCs (**Figure 2A**). To further increase stringency, we defined differential methylation as a > ± 10% change between SA and control, yielding 126,456 differentially methylated sites (DMSs; **Figure 2B**, blue shaded columns), of which, 65.52% were hypomethylated and 34.48% were hypermethylated following cocaine SA suggesting a hypomethylation bias of cocaine SA.

**Figure 2.**
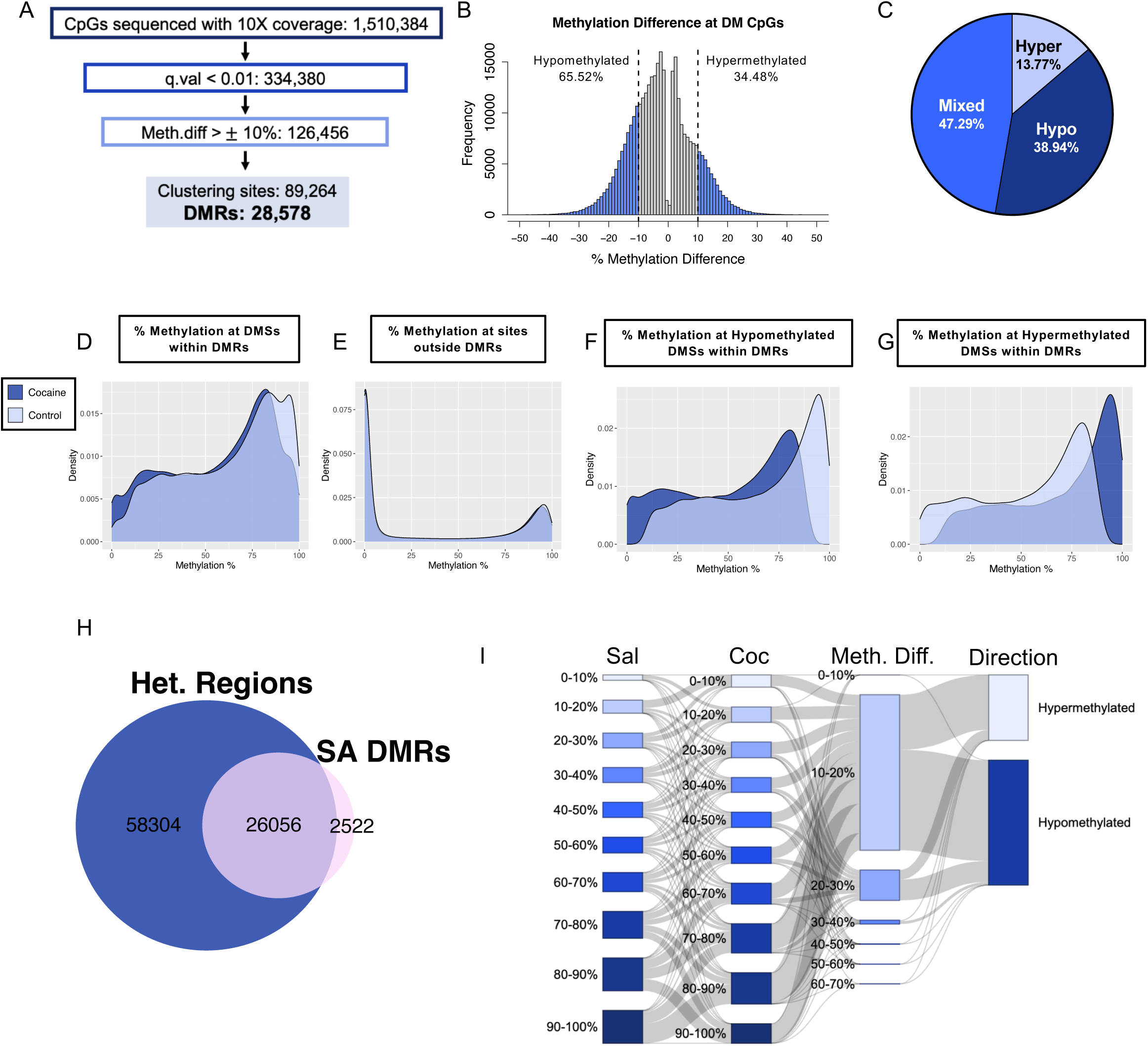
Large-scale DNA methylation changes in the dentate gyrus following cocaine SA. (**A**) Summary of eRRBS results of DNA methylation at CGs. Genomic DNA from 4 cocaine SA and 4 saline control animals was used for individual replicates in eRRBS. (**B**) Distribution of methylation differences (cocaine minus saline) of CGs with 10X coverage and q-value ≤ 0.01. Blue columns represent CGs with ≥10% methylation changes. (**C**) Distribution of SA-DMRs that contain all hypermethylated sites (13.77%), all hypomethylated sites (38.94%), or both hypermethylated and hypomethylated sites (47.29%). (**D**) Methylation percent distribution of clustered differentially methylated sites (DMSs) within SA-DMRs for saline (light blue) and cocaine (dark blue) groups. (**E**) Methylation percent distribution of CGs outside of SA-DMRs exhibits a bimodal distribution. (**F, G**) Methylation percent distribution of clustering (**F**) hypomethylated or (**G**) hypermethylated DMSs within SA-DMRs for saline (light blue) and cocaine (dark blue) groups. (**H**) Overlap between genomic regions with methylation heterogeneity (Het., ≥2 CGs with 10-90% methylation) in control DGCs and SA-DMRs. (**I**) Sankey diagram of distribution of percent methylation of DMSs for the saline group, the cocaine group, the methylation difference between the saline and cocaine groups, and the direction (hypo or hyper) of this methylation difference.

Next, we clustered 2 or more neighboring DMSs within 1kb of each other (**Supplementary Table 1**) into differentially methylated regions (SA-DMRs) between SA and control DGCs (**Supplementary Table 2**). On average, SA-DMRs spanned 227.9 base pairs and contained 3.12 DMSs (**Supplementary Figure 2**). We identified 28,578 SA-DMRs (**Figure 2A)**, representing 5.9% of all profiled CpGs, demonstrating extensive cocaine-induced reorganization of the DGC methylome. Similar to DMSs, a greater proportion of SA-DMRs contained solely hypomethylated CpG sites (hypo DMRs, 38.94%), compared to DMRs with solely hypermethylated CpG sites (hyper DMRs, 13.77%; **Figure 2C**). The remaining 47.99% were mixed DMRs, containing both hyper– and hypomethylated CpG sites.

### Cocaine SA targets genomic regions with intermediate methylation

Although the vast majority of CpGs in the genome are either uniformly methylated (∼100%) or unmethylated (∼0%) in homogeneous cell populations^36, 37^ (i.e., bimodal methylation pattern), DMSs within SA-DMRs exhibited methylation in the intermediate range (10% to 90%) in saline controls, indicating methylation bistability, i.e., the presence of both the methylated and unmethylated epialleles of each specific DMS across DGCs (**Figure 2D**, light blue). Since most reads in bulk eRRBS correspond to individual cells, it follows that SA-DMSs were in the methylated state in some while in the unmethylated state in other DGCs. CpGs outside of SA-DMRs, displayed genome typical bimodal methylation in yoked saline controls (**Figure 2E**). We have recently found that similar intermediately methylated regions were preferentially targeted for hypo-and hypermethylation during extinction learning, chronic wheel running, and repeated unpredictable stress in dorsal DGCs (unpublished data), as well as in fetal ventral DGCs by proinflammatory gestational environment^38^ suggesting that regions with methylation bistability are epigenetically highly malleable by external stimuli. However, SA altered methylation at ∼30,000 SA-DMRs, an unusually large number compared to the few thousand regions affected by non-drug environments. We propose that cocaine, by increasing dopamine and/or norepinephrine levels^39^, amplifies the entorhinal glutamatergic input onto DGCs and perturbs the native balance between the methylated and unmethylated states at a large number of methylation bistable regions.

Given the large number of methylation bistable regions that are sensitive to and epigenetically malleable by cocaine SA (**Figure 2A**), we next explored the total number of regions with similarly clustered, intermediately methylated (10-90%) CpGs in the DGC genome of yoked control animals. We identified 84,360 genomic blocks containing at least two clustered intermediately methylated CpGs, with an average block size of 275.8 bp, which is comparable to the 227.9 bp length of SA-DMRs. Nearly all (91.18%) of the SA-DMRs overlapped with these genomic blocks (**Figure 2H**), further confirming that cocaine SA preferentially targets sites/regions with an intermediate methylation pattern. Given that ∼1/3^rd^ of all methylation bistable regions were modified by cocaine, we concluded that cocaine SA is associated with a large-scale reorganization of methylation at bistable regions in the mouse dorsal dentate gyrus.

### Methylation state switching involves only a subset of dentate granule cells

Cocaine SA shifted the methylation by an average of 15.70% (10-30%) to a lower level at hypomethylated DMR DMSs (**Figure 2B, F**), thereby increasing the proportion of the unmethylated epiallele of these DMSs/DMRs and the fraction of cells with the unmethylated epialleles in the DGC population. Conversely, the smaller number of hypermethylated DMSs in SA-DMRs exhibited the opposite trend with an average of 16.16% (10-30%) increase in methylation, shifting the fraction of cells with the methylated epialleles to higher (**Figure 2B, G**). These trends in epiallelic switching were most apparent when DMSs and their cocaine-induced methylation changes were analyzed in 10% methylation bins (**Figure 2I**). Whether these changes occur in overlapping sets of DGCs is unknown. However, given that experiences in the dentate gyrus are encoded in small neuronal ensembles (estimated to be 7-8% by calcium imaging^40^) and that cocaine SA unfolds over multiple days, it is reasonable to hypothesize that most epiallelic switching reflects a cumulative population of DGCs activated throughout SA. Higher methylation shifts at DMSs could correspond to the cell population that was repeatedly recruited, while lower methylation shifts may be related to cell populations that entered or left the ensemble during SA or may represent DMSs that were less amenable for methylation switching.

### Cocaine-induced differentially methylated regions are enriched in transcriptional enhancers

Heterogeneity in DNA methylation is a defining feature of cis-regulatory elements, particularly of enhancers^41–45^, which play a key role in gene regulation. To explore if enhancers are overrepresented in SA-DMRs, we next examined their chromatin state in control DGCs using annotations of chromatin states as defined by ChromHMM^46^ in C57BL6/J mouse DGCs^47^. ChromHMM categorizes chromatin into distinct states based on the combinatorial presence and absence of histone modifications, including promoter, enhancer, and heterochromatin states. Of the 28,578 SA-DMRs identified, 50% were in cis-regulatory elements (**Figure 3A**). These regions were enriched in both active (A) enhancer (Odds Ratio: 1.555, FDR = 4.40E-16) and poised (P) enhancer (Odds Ratio: 1.230, 4.40E-16) chromatin signatures, based on the distinct combinations of the histone marks H3K4me1, H3K4me3, and H3K27ac (**Figure 3B**). Enhancer SA-DMRs had a comparable distribution of hyper, hypo, and mixed DMRs (**Supplementary Figure 3**) as observed for all the SA-DMRs (**Figure 2C**).

**Figure 3.**
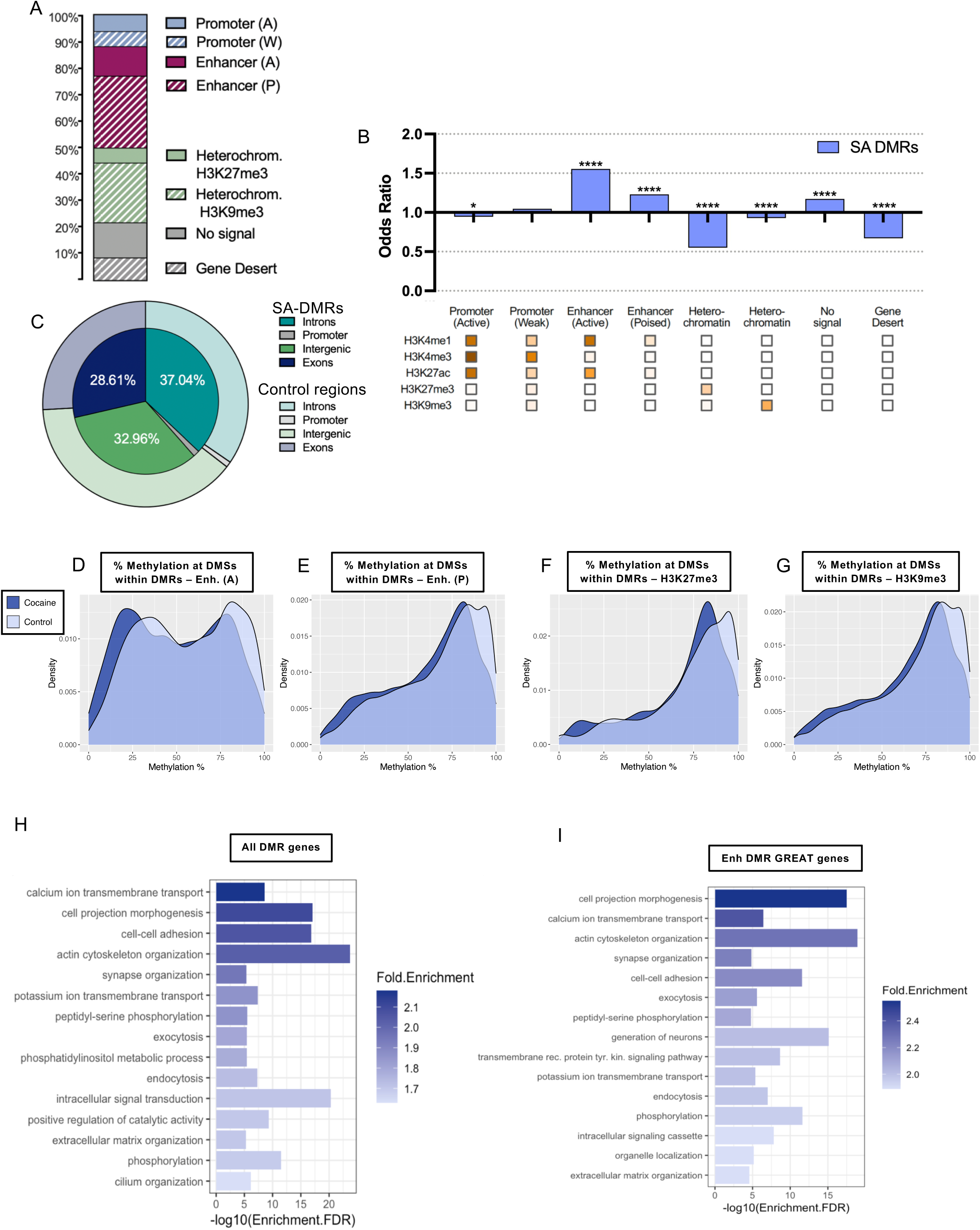
Enrichment of differentially methylated regions in enhancers and genes. (**A**) Distribution of SA-DMRs across eight chromatin states as determined by chromHMM^46^ in mouse DGCs^47^. (**B**) SA-DMRs are enriched, based on odds ratio, in active (Odds Ratio: 1.555, FDR = 4.40E-16) and poised (Odds Ratio: 1.230, FDR = 4.40E-16) enhancers. The heatmap (bottom), adapted from Zhu et al.^47^, shows the emission probability of relevant histone marks identified by chromHMM^46^. * p-value ≤ 0.05, **** p-value ≤ 0.0001 (**C**) Distribution of SA-DMRs across gene features as compared to randomized control regions. (**D-G**) Methylation percent distribution of clustering DMSs within SA-DMRs corresponding to four chromatin signatures for saline (light blue) and cocaine (dark blue) groups. (**H**) Gene ontology^89, 90^ enrichment of genes containing SA-DMRs in biological processes. **(I**) Same as H for genes associated with enhancer SA-DMRs, as determined by the GREAT database^51^.

Although ∼30% of SA-DMRs were in facultative H3K27me3 and constitutive H3K9me3 heterochromatin states (**Figure 3A**), SA-DMRs were relatively depleted in these chromatin features because of the abundance and wide distribution of heterochromatin in the genome (**Figure 3B**). Further analysis revealed that about two-thirds of SA-DMRs were located within the gene body, made up of exons and introns (**Figure 3C**, inner circle). This was a significant enrichment when compared to the distribution of randomized control regions (**Figure 3C**, outer circle; Odds Ratio: 1.252, FDR = 3.67E-16). In contrast, SA-DMRs were depleted in intergenic regions as compared to the control regions (Odds Ratio: 0.677, FDR = 3.67E-16), suggesting that the majority of SA-DMRs, including those associated with enhancer signatures, are concentrated in protein-coding regions of the mouse genome rather than in intergenic regions.

Given the abundance and enrichment of SA-DMRs in enhancer specific chromatin states in control DGCs (**Figure 3A**), we next examined baseline methylation and cocaine induced methylation shifts specifically in the group of enhancer-associated SA-DMRs. Similar to all SA-DMRs (**Figure 2D**), active enhancer-associated SA-DMRs exhibited intermediate methylation, although methylation extended toward the lower intermediately methylated range (**Figure 3D**). This pattern is consistent with the generally low methylation of active enhancers across various cell types and tissues^43, 48^. In contrast, SA-DMRs located in poised enhancers displayed higher average methylation levels (**Figure 3E**), similar to the pattern of heterochromatin associated DMRs (**Figure 3F, G**), a pattern consistent with their putative inactive chromatin state.

### Cocaine-induced differentially methylated regions are associated with a large number of neuronal genes

Given the unusually large scale of cocaine SA-induced methylation changes (**Figure 2I**) and their concentration in gene bodies (**Figure 3C**), it was not surprising that the 28,578 SA-DMRs mapped to 9,833 genes. The large number of these DMR genes (SA-DMGs) suggested that functionally diverse genes harbor one or more regions that are epigenetically responsive to cocaine SA. However, DMGs were overrepresented in specific Gene Ontology (GO) biological processes including “cell adhesion,” “calcium ion transport,” “extracellular matrix,” and “phosphorylation” among others (**Figure 3H**). While not all these processes are strictly neuron-specific, they all have relevance to neuronal processes. Remarkably, SA-DMGs were not enriched in general cellular processes such as transcription, transcription factors, RNA binding proteins, translation, and energy metabolism (FDR >0.05), categories that are typically represented by a large number of genes^49, 50^. This suggests that cocaine SA-induced methylation changes are not randomly distributed across the DGC genome and genes but are selectively concentrated in genes linked to neuronal function.

Next, we focused our analysis on DMGs associated with active and poised enhancer SA-DMRs because these cis-regulatory elements were overrepresentation in DMRs (**Figure 3B**). Enhancer-specific SA-DMRs could be linked to 6,995 genes as determined by the GREAT database^51^, which connects cis-regulatory elements to their target genes. Enhancer DMR DMGs were enriched in many of the same GO biological processes as the larger group of all DMGs (**Figure 3I**). The highly similar gene sets associated with enhancer and all SA-DMRs was largely due to the many genes that contained both enhancer and non-enhancer DMRs.

### Genes differentially expressed following cocaine self-administration are enriched in cis-regulatory DMRs

As epigenetic modifications of cis-regulatory elements have downstream effects on gene expression, we performed bulk RNA-sequencing on DGCs isolated from the dentate gyrus of mice after cocaine SA (**Figure 1A**). Analysis of differential expression in 4 cocaine SA mice and 5 control mice by PCA clearly showed the separation of the two groups (**Supplementary Figure 4**). We found 361 upregulated and only 23 downregulated cocaine SA genes (**Figure 4A**). Of these 384 differentially expressed genes (SA-DEGs, **Supplementary Table 3**), 184 contained SA-DMRs (i.e., were SA-DEG x DMGs), representing a significant enrichment (hypergeometric testing: p-value < 0.02; **Figure 4B**). Nevertheless, many more SA-DMR genes did not show expression differences between cocaine SA and yoked control animals, suggesting that only a specific set of DMRs and/or DMGs may be associated with differential expression following cocaine SA.

**Figure 4.**
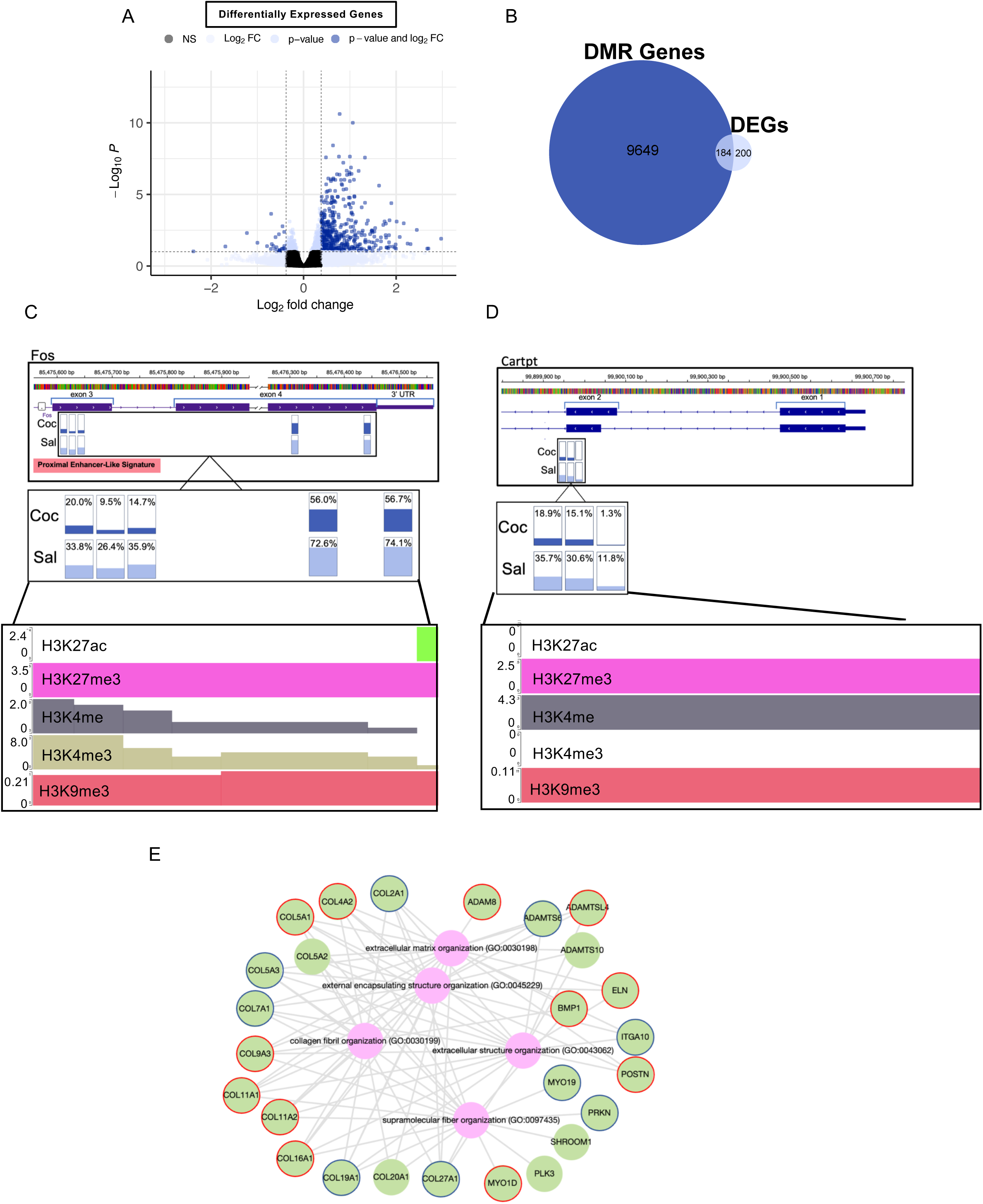
Upregulation of regulatory genes and extracellular matrix-related genes in SA DGCs. (**A**) Volcano plot of differentially expressed genes after seven days of cocaine SA, (black: non-significant by p-value and below log2FC threshold; light blue: significant by adjusted p-value or above log2FC threshold, but not both; dark blue: significant (adjusted p-value < 0.1) and above log2FC cutoff of log2(1.3). RNA-Seq with 4 individual cocaine SA and 5 individual control mice. (**B**) Overlap of differentially expressed and differentially methylated genes (DEGs x DMGs). Hypergeometric test, p-value < 0.02. (**C**) The regulatory DEG x DMG c-*fos* with associated DMR (inset) spanning exons 3 and 4 and containing 5 DMSs (represented by vertical rectangles). DMSs are hypomethylated in cocaine as compared to saline. Light and dark blue fills represent percent methylation in saline and cocaine groups, respectively (Gene view from IGV^91^). In the magnified image, % methylation is indicated. Lower panel shows autoscaled chromatin marks in *c-fos* DMR as defined by ChromHMM in DGCs. The region has both activating (H3K4me1 and H3K4me3) and silencing (H3K27me3, H3K0me3) marks. (**D**) Same as “C” depicting the DMR (inset) associated with the regulatory DEG x DMG *cartpt,* containing 3 DMSs that are hypomethylated in cocaine as compared to saline. This region is associated with activating (H3K4me1) and silencing (H3K27me3, H3K9me3) marks. (**E)** Enrichment of the 361 upregulated DEGs in GO biological processes (created with enrichr-KG^92^). Genes associated with enhancer SA-DMRs are outlined in red. Genes containing non-enhancer SA-DMRs are outlined in navy.

We characterized SA-DMRs located within DEGs to determine whether they exhibited any unique features compared to all SA-DMRs. Regulatory elements, in particular active promoters and poised enhancers were represented by a higher relative proportion in DEG associated DMRs compared to the much larger group of all DMRs (63% vs 50%, **Supplementary Figure 5A** and **Figure 3A**). Also, DEG-associated DMRs had a higher enrichment score in cis-regulatory elements than DMRs (**Supplementary Figure 5B** and **Figure 3B**). DMRs in DEGs were somewhat larger than the average size of DMRs (291.8 bp vs. 227.9 bp, respectively) and contained slightly more DMSs (3.5 vs. 3.1, respectively). Otherwise, we found no difference in baseline methylation levels or range between DMSs in DEG associated DMRs and all DMRs (**Supplementary Figure 6** and **Figure 2D**), and the extent and direction of SA-induced methylation shifts were similar in the two groups (16-17% and 2:1 hypo– to hypermethylated ratio).

### Upregulation of genes with regulatory and extracellular matrix (ECM)-related functions following cocaine self-administration

Drugs of abuse reorganize gene regulatory networks (GRNs) in neuronal populations associated with reward and memory^52, 53^. GRNs are hierarchical regulatory frameworks organized into modules around hubs of regulators such as transcription factors (TFs) and signaling molecules that are highly connected to downstream genes. We identified two possible regulatory genes in cocaine SA, *c-fos* and *cartpt* (cocaine and amphetamine-regulated transcript prepropeptide) that were both upregulated and hypomethylated in SA DGCs (**Figure 4C, D**). FOS is a key TF in GRNs, acting as an immediate early gene that responds rapidly to stimuli, such as acute cocaine administration in various brain regions, including the dentate gyrus^54–56^. Hypomethylation of the putative enhancer-like region in the proximal part of the DMR (**Figure 4C**) suggests a more persistent upregulation of *c-fos* by cocaine SA. FOS with JUN family members forms the AP-1 (Activator Protein 1) complex that activates numerous downstream genes involved in many functions, including hippocampal learning and memory^57^. The DMR in *c-fos* had both activating (H3K4me1 and H3K4me3) and silencing (H3K9me3 and H327me3) marks in DGCs (**Figure 4C**, lower panel), indicating bivalency^58^ at the *c-fos* DMR. Theoretical models suggest that co-occurrence of active and repressive chromatin marks on the same nucleosome may represent the transition point between active and silent states^59^. Such chromatin bistability would align with the methylation bistability of the *c-fos* DMR.

*Cartpt*, the second putative regulatory gene hypomethylated and upregulated by cocaine SA, encodes a signaling peptide (CART) that, by interacting with GPR160, leads to the activation and phosphorylation of ERK and the transcription factor CREB (cAMP response element-binding protein^60, 61^. Although it is activated by acute amphetamine and cocaine^62, 63^, its hypomethylation argues for a more persistent upregulation following cocaine SA (**Figure 4D**). The *cartpt* DMR was also associated with both activating and silencing chromatin marks (H3K4me1, H3K27me3, and H3K27me3), in line with its methylation bistability (**Figure 4D**, lower panel).

Next, we performed GO analysis with all SA-DEGs that showed an overrepresentation in the biological process of “ECM organization” as well as other ECM-associated categories (**Figure 4E**). This GO term was also among the top 15 biological processes enriched in DMGs (**Figure 3H, I**). The ECM is a three-dimensional molecular network surrounding cells that is essential for neuronal plasticity^64, 65^. All 26 ECM-related DEGs were upregulated, including many collagen genes; 12 of them were also genes associated with enhancer SA-DMRs (**Figure 4E**, outlined in red) and 9 more contained non-enhancer SA-DMRs (**Figure 4E**, outlined in blue). Although the hypomethylation bias in SA-DMRs is consistent with the cocaine-induced upregulation of ECM genes, the methylation pattern of these DMRs was complex, with some containing both hypo– and hypermethylated DMSs. The other GO functional categories, enriched in SA-DMGs (**Figure 3H**), were not represented at a significant level by DEGs, suggesting that ECM genes are unique among the SA-DMR genes, and could represent a co-regulated GRN module involved in cocaine-induced hippocampal plasticity and cocaine SA behavior.

## Discussion

This work underscores the importance of considering the role of the dentate gyrus in cocaine-related behaviors, even in tasks traditionally associated with cue processing, such as cocaine SA. Our data demonstrates that cocaine SA produces DNA methylation changes in DGCs across a large number of small genomic regions, exhibiting heterogenous methylation. This methylation heterogeneity reflects the coexistence of methylated and unmethylated epialleles within the population of genetically identical and morphologically indistinguishable DGCs. Although methylation at individual DMSs in both saline control and SA animals was variable, their methylation levels across individuals within the saline control and within the SA groups were highly correlated (i.e., within group variation, r^2^ = 0.808 ± 0.015 and 0.793 ± 0.036, respectively). Correlation between pairs of individual SA and control animals was lower (i.e., between group variation; 0.723±0.029) indicating that heterogeneity in baseline DMS methylation was not due to individual variability but arose stochastically (**Supplementary Figure 7**).

Cocaine SA was associated with a 10-30% methylation shifts at DMSs. Because of the predominance of hypomethylation over hypermethylation, cocaine SA primarily resulted in an increase in the unmethylated epialleles at the expense of the methylated epialleles. A similar hypomethylated bias was reported in the NAc after cocaine SA^6^, and pharmacological inhibition of DNA methyltransferase activity, which decreases methylation, increased cocaine SA^5^, suggesting a causal relationship between drug-induced DNA hypomethylation and drug taking.

The 10-30% (mean ∼16%) methylation shifts induced by cocaine SA indicate epiallelic switching at DMSs/DMRs in a similar fraction of DGCs. Due to sparse coding in the dentate gyrus^67, 68^, only a small fraction of DGCs, estimated to be a few % by two-photon calcium imaging^40^, is activated in a given environment. DGCs have a relatively stable spatial code over days^40^, but ensembles are dynamic with new cells entering and previously activated cells exiting the ensemble^69^. Therefore, it is reasonable to suggest that the overall size of the SA ensemble is increased through the 7-day SA. Since DMRs are likely more amenable to methylation changes in activated than in silent neurons, we posit that neurons activated through SA are those that undergo methylation switching.

The large number of SA-DMRs was striking, especially given that non-drug associated, and purely environmental challenges produced an order of magnitude fewer DMRs in the dentate gyrus. Consistent with the high number of DMRs, a large number of genes were associated with DMRs, and these DMGs, as expected, were enriched in diverse GO functional categories. However, all DMGs had relevance to neuronal functions, rather than being randomly distributed among genes. The large number of SA-DMRs and DMGs likely reflects the increased levels of dopamine and/or norepinephrine during cocaine SA that may amplify the entorhinal glutamatergic input onto DGCs. Indeed, DRD1 in the hippocampus has been shown to be essential for the formation of cocaine-paired contextual memory and cocaine seeking after SA^39^. Moreover, long-term potentiation (LTP), a proxy for hippocampal learning, was reduced by DRD1/5 antagonists, while L-DOPA and a DRD1/D5 agonist reduced the threshold for LTP induction^70^. Similarly, cocaine was reported to enhance LTP in the CA1 area of the hippocampus^71, 72^.

In contrast to the many thousands of SA-DMGs, only ∼400 genes were differentially expressed in SA DGCs, half of which were also differentially methylated. While the number of DEGs containing SA-DMRs exceeds chance expectations, the large number of DMGs with no expression difference in bulk RNA-seq suggests that methylation changes at specific sets of DMRs/genes may produce detectable differential expression. Indeed, genes are highly interconnected in hierarchical GRNs and the gene expression output of GRNs is robust against perturbation, with master regulators and their downstream target modules being the most responsive to changes in the environment^73^. We identified two groups of differentially methylated and differentially expressed genes, both with functional relevance to cocaine-induced neuroplasticity and cocaine SA. First, *c-fos* and *cartpt*, two regulatory genes that are induced by cocaine^55, 62, 63, 74^ and are potential upstream regulators of cocaine-induced neuroplasticity and SA^6, 75^ were hypomethylated and upregulated in SA-DGCs. Cocaine administration was shown to increase *c-fos* expression^55, 74^, and its promoter was reported to be hypomethylated in the NAc and mPFC following reinstatement of cocaine SA, a change that could be reversed by methyl supplementation^6^. In our cocaine SA protocol, upregulation of *c-fos* in SA DGCs was accompanied by hypomethylation at a putative enhancer in exon 3. The other upregulated regulatory gene, *cartpt*, and its DMR also showed hypomethylation and association with an enhancer specific histone mark. *Cartpt* is upregulated after stimulant use^62, 63, 76^ and its protein, CART, exerts a positive neuromodulatory effect on psychostimulant SA. CART activates the ERK signaling cascade through NMDA receptors in the hippocampus, potentially connecting CART expression to contextual learning^77^. Furthermore, pharmacological and genetic inactivation of DRD1 blocks the cocaine-induced increase of CART expression^75^, suggesting that upregulation of *cartpt* in SA DGCs may be due to the cocaine-induced increase in dopamine and subsequent activation of DRD1 in DGCs.

The second group of SA-DEGs consisted of a significant number of ECM-associated genes. The ECM plays a key role in synaptic plasticity as it modulates both NMDA receptors and L-type voltage-gated calcium channels, important players in cocaine’s effects^78, 79^, to trigger activity-dependent changes^64^. Moreover, ECM-related gene expression was shown to be modulated by cocaine use in hippocampal tissues from humans and nonhuman primates^80, 81^, suggesting a link to drug use and addiction. Within the ECM cluster, we identified 10 differentially methylated and expressed collagen genes, which is notable because a similar type of contextual learning, contextual fear conditioning, was reported to alter expression of collagen genes in the CA1 region^82^. Additionally, heroin SA has been associated with changes in ECM-related genes, including collagen genes, in brain regions associated with reward learning such as the basolateral amygdala, ventral hippocampus, and ventral tegmental area^84^. Two of the upregulated and differentially methylated collagen genes in our experiments, *Col7a1* and *Col27a1*, were also upregulated in the NAc after cocaine conditioned place preference learning^85^.

Although no GRN model is available for cocaine SA to connect regulatory genes with their targets and ultimately to the cellular and behavioral phenotypes, FOS proteins form AP-1 complexes with JUN proteins, which bind to DNA and control the transcription of many genes, including those encoding ECM components, albeit this link has only been established in connective tissue^86, 87^. CARTPT does not directly activate ECM genes but plays roles in regulating tissue remodeling, especially in bone and blood vessels, by influencing related pathways, leading to changes in ECM gene expression^88^.

In summary, we have demonstrated that cocaine SA is associated with large-scale DNA methylation changes and altered expression of specific sets of genes in the dorsal dentate gyrus of adult male mice. The methylation changes were predominantly hypomethylation and occurred in enhancers, supporting a link between drug-induced methylation changes, in particular hypomethylation, and cocaine-induced hippocampal plasticity and SA behavior. The large-scale effect of cocaine on the DGC methylome could be due to the convergence of contextual information from the entorhinal cortex and cocaine-enhanced dopamine and norepinephrine signaling in the dentate gyrus. A specific group of genes encoding ECM proteins, previously implicated in hippocampal structural plasticity and learning, emerged as both differentially methylated and upregulated following cocaine SA. The expression changes may contribute to drug-associated neuroplasticity in DGCs and to the contextual memory associated with cocaine SA.

## Acknowledgements

We would like to thank the support of the Epigenomics Core and the Genomics Core of Weill Cornell Medicine.

## Author contribution

Contributed to the conception of the work: MRB, EB, AR, MT Drafted the work or revised content: MRB, KAW, RS, AR, MT Approval of the version: all authors Accountability: all authors

