## Supplemental Figures 1-7 for "Large-scale reorganization of DNA methylation and upregulation of extracellular matrix genes in the dorsal dentate gyrus following cocaine taking"

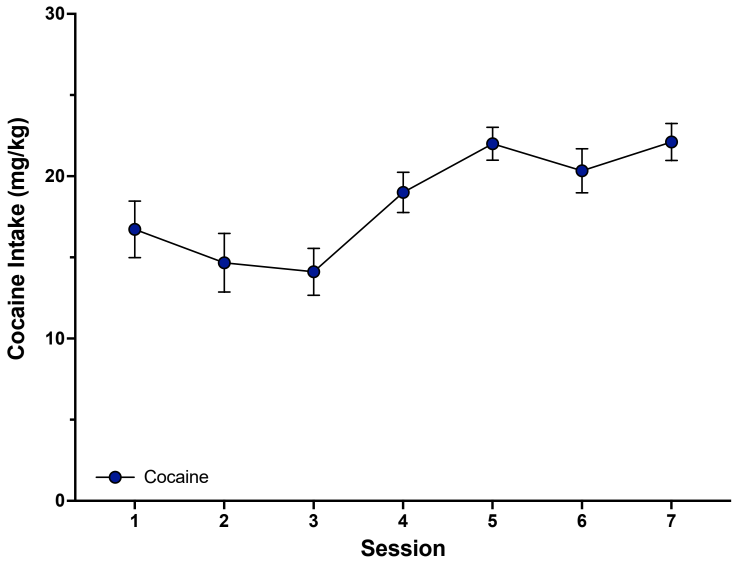


**Supplementary Figure 1. Cocaine intake through seven sessions of cocaine SA**. Significant increase in intake of cocaine through the sessions, one way repeated measures ANOVA, $F(2.43, 14.58) = 4.767$, $P=0.0207$.

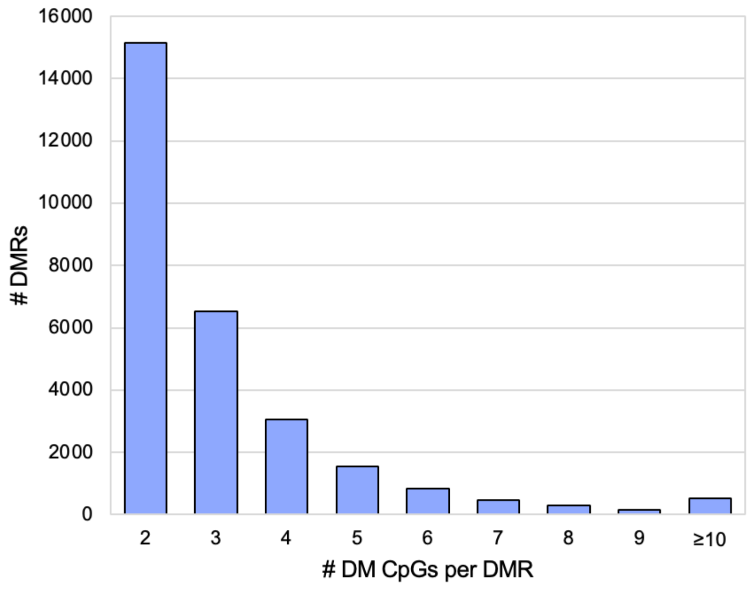


**Supplementary Figure 2. Distribution of SA-DMRs based on their DMS content.**

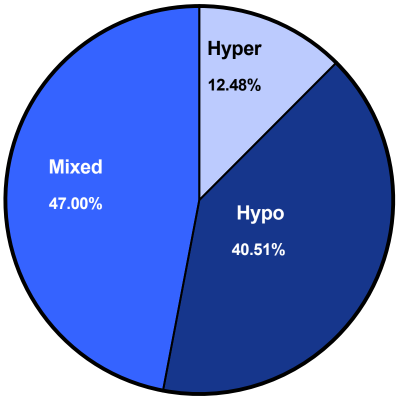


**Supplementary Figure 3.** Distribution of enhancer-associated SA-DMRs that contain all hypermethylated sites, all hypomethylated sites, or both hypermethylated and hypomethylated sites.

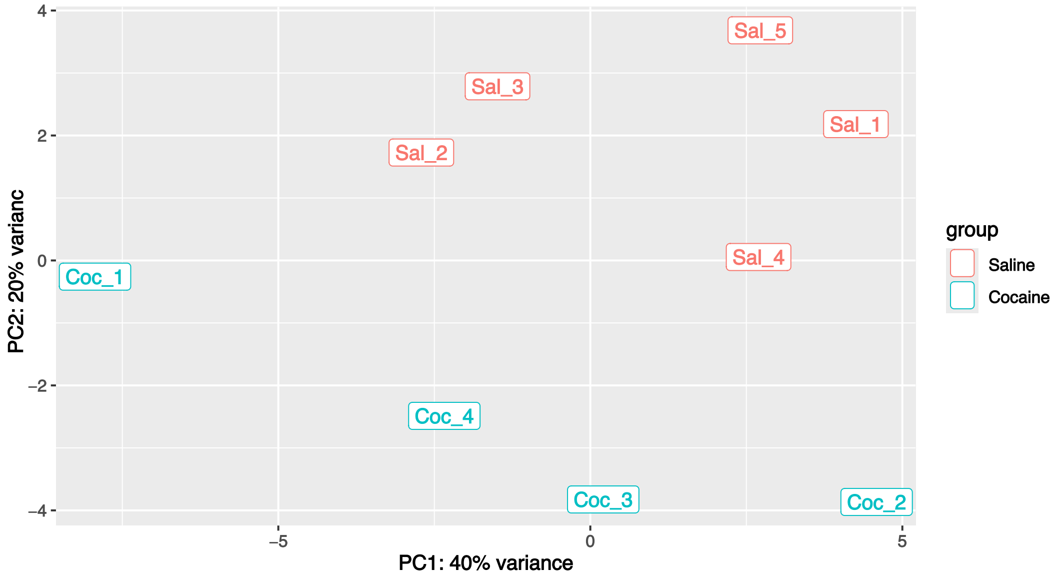


**Supplementary Figure 4. PCA plot of dentate granule cell RNA-seq data from individual saline control (Sal) and cocaine SA (Coc) mice.** A combination PC1 and PC2 explains most  variance between cocaine SA (Coc) and Saline control (Sal) samples.

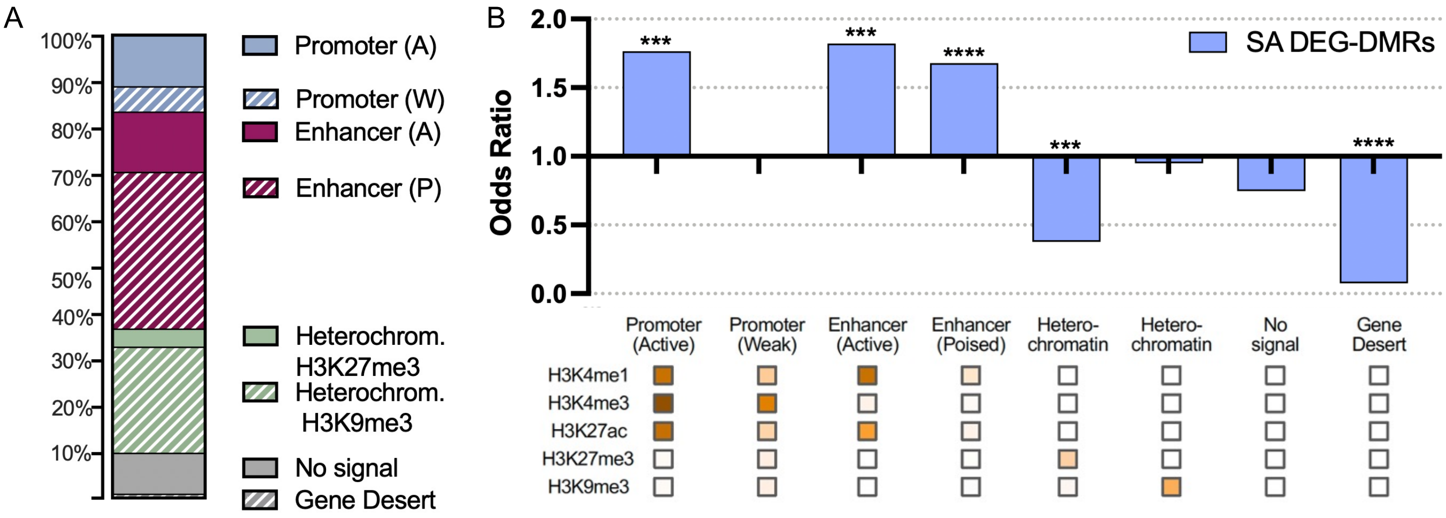


**Supplementary Figure 5. Chromatin states of SA-DMRs. A.** Distribution of DEG-DMRs across eight chromatin states as determined by chromHMM in mouse DGCs. **B.** Enrichment of DEG-DMRs in chromatin states, *** p-value < 0.001, **** p-value < 0.0001.

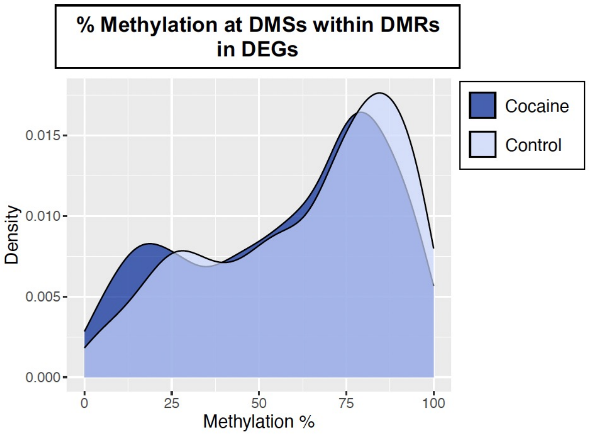


**Supplementary Figure 6. Percent methylation at DMSs within DEG-DMRs.**

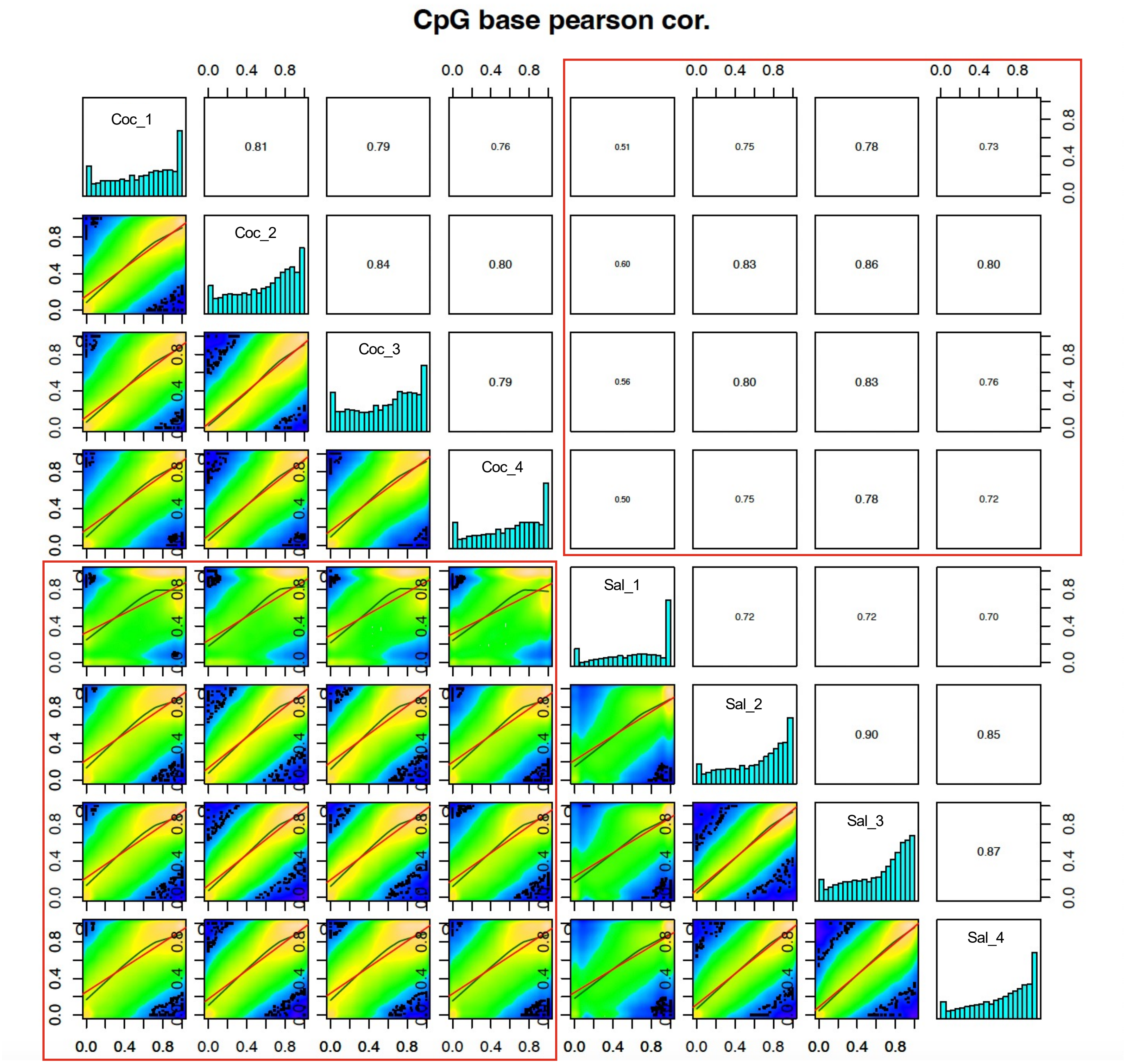


**Supplementary Figure 7. Higher correlations in DMS methylation within the group of cocaine SA animals ( Coc; r2 = 0.793 + 0.036, mean + SE) and saline controls (Sal; 0.808 ± 0.015) than between individual saline and cocaine SA animals (0.723+0.029).** Correlations between pairs of cocaine SA and control animals are highlighted in red boxes. The more deviation of cocaine SA vs. saline correlations from the trendline (in the lower left red box), especially at the higher methylation range, reflects the hypermethylation bias in differential methylation by SA.
