## Supplementary Methods for "Large-scale reorganization of DNA methylation and upregulation of extracellular matrix genes in the dorsal dentate gyrus following cocaine taking"

### Animals

Animal experiments were performed in accordance with The Rockefeller University and Weill Cornell Medicine Institutional Animal Care and Use Committee guidelines. Adult C57BL/6 male mice (11 weeks upon arrival; Jackson Laboratory, Bar Harbor, ME) were group housed in a climate-controlled environment, 4-5 animals per cage, with a reverse 12 h light/dark cycle (7PM-7AM). Food and water were available *ad libitum*.

### Surgery

Following one-week habituation, mice underwent jugular catheterization surgery as described previously[31, 32]. Briefly, animals were anesthetized with isoflurane and an indwelling catheter (0.012" ID by 0.025" OD) inserted into right jugular vein. Animals were allowed to recover for 5-7 days prior to the beginning of the operant SA procedure.

### Intravenous self-administration

Following recovery from surgery, mice had 7 consecutive daily 2-hour cocaine intravenous SA (n=12) or yoked saline (n=11) sessions in standard mouse Med Associates operant chambers equipped with two nose poke manipulanda (ENV-307A; St. Albans, VT). Animals were randomly allocated to the SA and yoked saline groups. For cocaine SA, mice were trained to nose poke at the active hole of the chamber for delivery of 0.5 mg/kg infusion of cocaine for 7 days on a fixed ratio 1 (FR1) reinforcement schedule with a 20 second "time-out" period following each infusion. Responding on the inactive nose poke was recorded but had no programmed consequence. Acquisition criteria included at least 10 infusions per session and a minimum of 70% of responses on the active versus inactive nose poke (indicating that the mouse learned the association of active nose poke and cocaine infusion). For the yoked saline controls, animals received an equivalent infusion of saline when their yoked paired cocaine SA mouse received an infusion of cocaine. The two nose poke holes were differentiated as the "left side" and "right side". Nose poke responses were recorded but had no programmed consequence. Following the seventh session, catheter patency was accessed via low dose intravenous ketamine. Three cocaine SA mice were excluded from analysis (two lost catheter patency and one failed to self-administer). Four randomly chosen mice per group were used for eRRBS and five separate mice per group were used for RNA-sequencing.

**DNA extractions for eRRBS (enhanced reduced representation bisulfite sequencing)**

Brains from adult mice were collected, flash frozen on dry ice, and stored at -80$^{o}$C. Brains were sectioned into 200 μm slices in a cryostat. The DGC cell bodies of the dorsal dentate gyrus (i.e., granule cell layer) were micro-dissected. DNA was isolated using the QIAamp DNA Micro Kit (QIAGEN) according to manufacturer instructions.

**Bisulfite Sequencing by eRRBS**

Genomic DNA from 4 cocaine SA and 4 saline control animals was used for individual replicates in eRRBS. Library preparations, sequencing, and adapter trimming was performed by the Epigenomics Core at Weill Cornell Medicine as described previously[33]. Single end 100 bp eRRBS sequencing, with a target of 100 million reads per sample, was performed on an Illumina NovaSeq6000 according to the manufacturer's instructions.

**Differential methylation analysis**

Alignment to the mm10 reference genome and methylation calling was performed using Bismark v22.1[34] using default settings. MethylKit v1.26[35] was used to perform differential methylation and statistical analyses in R v4.3.1. CpGs with at least 10 coverage reads were included. The coordinate files of the 8 subjects (4 cocaine SA and 4 saline controls) were merged and only

CpGs sequenced in all samples were included for differential methylation analysis. As described previously[36], differentially methylated sites (DMSs) were defined as sites with a 10% difference in methylation between the two groups with a sliding linear model (SLIM)-corrected p-values (q-values) of ≤ 0.01. Differentially methylated regions (DMRs) were defined as regions with two or more DMSs, with a DMS-to-DMS distance across all DMSs less than 1kb. DMRs were classified as hypomethylated, hypermethylated, or mixed based on whether all DMSs in the DMR were hypomethylated, hypermethylated, or a combination of the two.

**Enrichment of DMRs in annotated chromatin states and genomic features**

We analyzed enrichment of DMRs in eight chromatin states previously determined by chromHMM[37] for mouse DGCs[38]. For enrichment of DMRs in genomic features we used exon, introns, 3'UTR, 5'UTR, and TSS annotations from the USCS genome browser[39] based on the mm10 genome. Promoters were defined as regions ± 500 bp from the transcription start site (TSS). To evaluate enrichment or depletion of DMRs in a specific chromatin or genomic feature, we calculated the Odds Ratio [(actual)/(potential – actual)]. Statistically significant enrichments were computed using Chi-square test in R, and p-values were adjusted using the Benjamini-Hochberg method. Control regions were created by clustering non-DMS CpGs sequenced in the genome into regions fitting the DMR definition of at least 2 CpG sites within 1kb.

**Assignment of DMRs to genes**

DMRs were assigned to their host genes through locating the DMRs within mm10 genome coordinates associated with genes. Enhancer DMRs were assigned to their potentially regulated genes by the GREAT database[40].

**Motif enrichment analysis**

We performed motif discovery with DMRs extended ±4 bp using SEA[41] with default settings in MEME Suite 5.5.1[42], with the most recent version of the HOCOMOCO (v11) mouse TF binding site collection[43]. Significance was determined as e-value ≤ 10 and q-value ≤ 0.05.

**RNA extractions**

Brains from 5 cocaine SA and 5 saline control mice were collected, flash frozen on dry ice, stored at -80°C, and cryostat sectioned into 200 µm slices. Then, the granule cell layer was micro-dissected from the sections and total RNA was isolated from individual animals using the RNeasy Mini Kit (Qiagen, Valencia, CA). RNA for bulk RNA-sequencing all had RIN value ≥ 9. Because simultaneous RNA and DNA isolation from microdissected tissue had a lower yield, DNA for eRRBS and RNA for RNA-Seq were obtained from different SA and control mice.

**Bulk RNA-sequencing**

Library preparation and sequencing was performed by the Weill Cornell Medicine Genomics Core using NEB Ultra II Directional RNA Library Prep (plus Poly A isolation module), according to manufacturer protocol. Paired-end RNA sequencing was performed on an Illumina NovaSeq 6000 machine with 2x50bp reads with samples sequenced on an SP flow cell. Sequencing reads were trimmed with cutadapt[44] and aligned to the mm10 reference genome using STAR by the Weill Cornell Medicine Genomics core. Abundance of transcripts was measured with Cufflinks[45] in Fragments per Kilobase of exon model per million mapped reads (FPKM). Read counts were counted using the HT-seq program[46].

**Differential expression analysis**

Differential expression analysis was done using DESeq2 package in R[47]. Genes with less than 10 counts were pre-filtered out. Outlier detection was performed with robust principal component analysis, with the PCAGrid function[48]. Differentially expressed genes were determined using Benjamini-Hochberg corrected p-value < 0.1 threshold and log2FC > log2(1.3).

**Gene ontology and network analysis**

Panther GO-Slim overrepresentation analysis was performed using pantherdb.org. GO Biological Processes with <500 genes were used to identify specific (child) terms and avoid inflation of FDR values with large functional categories [49-51]. Network analysis was done using enrichr-KG[52].

**Statistical Analysis**

Statistical analyses for the cocaine self-administration behavior were performed in GraphPad Prism. Behavioral data had normal distribution as determined by the Shapiro-Wilk test. Behavioral data were analyzed with repeated measures two-way ANOVA (nose poke type (active versus inactive) x session) followed by Tukey multiple comparison posthoc test, when main effect was observed. Significance of overlaps between differentially expressed and differentially methylated genes was determined using hypergeometric testing (*phyper* in R). Statistics for eRRBS and RNA-Seq are incorporated into the corresponding method section. Sample sizes were based on previous work in the lab and are similar to other published work in the field. Sample sizes for individual experiments are indicated in figure legends.

**Code availability**

Standard packages were used for methylation and RNA-sequencing analysis as described in the specific results sections. Detailed code is available upon request.
